# Bi-directional allosteric pathway in NMDA receptor activation and modulation

**DOI:** 10.1101/2024.04.16.589813

**Authors:** Paula A. Bender, Subhajit Chakraborty, Ryan J. Durham, Vladimir Berka, Elisa Carrillo, Vasanthi Jayaraman

## Abstract

N-methyl-D-aspartate (NMDA) receptors are ionotropic glutamate receptors involved in learning and memory. NMDA receptors primarily comprise two GluN1 and two GluN2 subunits. The GluN2 subunit dictates biophysical receptor properties, including the extent of receptor activation and desensitization. GluN2A- and GluN2D-containing receptors represent two functional extremes. To uncover the conformational basis of their functional divergence, we utilized single-molecule fluorescence resonance energy transfer to probe the extracellular domains of these receptor subtypes under resting and ligand-bound conditions. We find that the conformational profile of the GluN2 amino-terminal domain correlates with the disparate functions of GluN2A- and GluN2D-containing receptors. Changes at the pre-transmembrane segments inversely correlate with those observed at the amino-terminal domain, confirming direct allosteric communication between these domains. Additionally, binding of a positive allosteric modulator at the transmembrane domain shifts the conformational profile of the amino-terminal domain towards the active state, revealing a bidirectional allosteric pathway between extracellular and transmembrane domains.

## Introduction

The majority of excitatory neurotransmission in the mammalian central nervous system is accounted for by glutamatergic signaling. The ionotropic glutamate receptor family includes N-methyl-D-aspartate (NMDA), α-amino-3-hydroxy-5-methyl-4-isoxazolepropionic acid (AMPA), kainate, and delta receptors^1, 2, 3, 4, 5, 6^. NMDA receptor signaling is involved in learning, memory, and synaptic plasticity, while dysfunction of NMDA receptors has been implicated in various neurological conditions, such as epilepsy, developmental delay, and ischemic stroke^7, 8, 9^. The NMDA receptor is a ligand-gated ion channel that requires simultaneous binding of glutamate and co-agonist glycine to activate^5, 10, 11^. NMDA receptors adopt a heterotetrameric structure, with each receptor comprising two obligatory glycine-binding GluN1 subunits and two glutamate-binding GluN2 (A-D) and/or glycine-binding GluN3 (A-B) subunits^3, 5, 12, 13, 14, 15^. These various subunit types share a conserved architecture consisting of extracellular amino-terminal (ATD) and agonist-binding domains (ABD), a transmembrane domain (TMD), and an intracellular carboxyl-terminal domain (CTD)^5, 12, 13, 16^. The binding of glutamate and glycine to the GluN2 and GluN1 agonist-binding domains induces conformational changes throughout the receptor, forming a transmembrane ion channel pore permeable to sodium, potassium, and calcium.

Several functional characteristics of NMDA receptors, including their extent of activation, desensitization profiles, and agonist potency, are determined by the identity of the GluN2 subunit^17, 18, 19, 20, 21^. GluN2A and GluN2D subunits represent opposite ends of the functional spectrum and differ in expression patterns. GluN1/GluN2A NMDA receptors are expressed broadly throughout the cortex and hippocampus^3, 17, 19^. In contrast, GluN1/GluN2D NMDA receptors show broad expression in early development but later become restricted to distinct subgroups of neurons, such as interneurons in the basal ganglia, hippocampus, and cerebellum^3, 17, 19^. Functionally, GluN1/GluN2A receptors demonstrate high open probabilities (∼50%), fast deactivation (∼50 ms), and lower glutamate potency (EC_50_ of 3-5 μM). In comparison, GluN1/GluN2D receptors have low open probabilities (< 4%), slow deactivation kinetics (> 1 s), and high glutamate potency (EC_50_ of 0.5 μM)^20, 22, 23, 24, 25, 26^. A chimeric study found that replacing the GluN2A amino-terminal domain with that of GluN2D decreased open probabilities compared to GluN2A wild-type receptors. In contrast, GluN2D receptor chimeras containing the GluN2A amino-terminal domain displayed increased open probabilities compared to wild-type GluN2D-containing receptors^22^. These findings suggest that the amino-terminal domain contributes to the functional and kinetic differences observed between NMDA receptor subtypes^22, 27^.

While several cryo-EM structures of the agonist-bound GluN1/GluN2A and GluN1/GluN2D receptors have been solved, the conformational differences underlying their differences in function could not be clearly identified^28, 29, 30, 31, 32^. Additionally, there are currently no apo state structures of the full-length GluN1/GluN2A or GluN1/GluN2D receptors. The lack of apo structures and unidentified significant conformational differences between the two subtypes is likely due to the highly dynamic nature of GluN1/GluN2D receptors. Recent single-molecule FRET (smFRET) investigations have determined the conformational landscape at the GluN1 amino-terminal domain^33^. However, recent structures of GluN2C-containing receptors reveal more significant conformational fluctuations across the amino-terminal domains of GluN2 subunits^28^. Hence, studying the conformational landscape at the GluN2 subunits in conjunction with protein dynamics is essential to gaining a comprehensive understanding of GluN2-dependent conformational variability underlying NMDA receptor subtype-specific gating properties.

The functional properties of NMDA receptors have been widely studied with respect to modulators^31, 34, 35, 36, 37, 38, 39, 40, 41, 42, 43^. Particularly, allosteric modulators have been observed to alter functional properties, such as open probability and potentiation. While extensive insight has been gained into how modulators that bind at the amino-terminal domain control channel activity, much less is known about modulators binding to the transmembrane segments^31, 34, 35, 36, 37, 38, 39, 40, 41, 42, 43^. The positive allosteric modulator GNE-9278 is particularly interesting as its binding site and functional properties have been well characterized^39^. GNE-9278 binds to the extracellular surface of the GluN1 pre-M1 transmembrane region and increases receptor currents in the presence of saturating concentrations of glutamate and glycine. Additionally, GNE-9278 slows deactivation kinetics and increases the potency of both glutamate and glycine. While GNE-9278 potentiates all GluN2-containing NMDA receptors, the most significant effect is seen for GluN1/GluN2D receptors with an approximately 8-to 15-fold increase in currents (7.9 for HEK cells, 14.9 for oocytes)^39^. While the binding site of this positive allosteric modulator has been determined, the mechanism underlying its function remains unclear.

Using single-molecule fluorescence resonance energy transfer (smFRET), we investigated the full spectrum of conformational states occupied by GluN1/GluN2D receptors in apo and agonist-bound conditions in comparison to GluN1/GluN2A receptors. We employed smFRET to measure distances across the GluN2 amino-terminal domain, the GluN1 and GluN2 agonist-binding domain clefts, and the GluN1 transmembrane domain. These measurements were used to categorize and compare conformational states and state transitions at the millisecond time scale. Our results reveal that the GluN2D amino-terminal domain is inherently more decoupled and dynamic in both apo and agonist-bound conditions. In contrast, the GluN1 transmembrane domain is more coupled and less dynamic in GluN1/GluN2D receptors than in GluN1/GluN2A receptors. The fraction of GluN1/GluN2D receptors coupled at the amino-terminal domain is correlated with the fraction of the receptors exhibiting longer distances at the transmembrane segments. Furthermore, this fraction also correlates with the low P_open_ of GluN1/GluN2D receptors, thereby providing a direct correlation between conformation and function of the receptor.

Additionally, we elucidated the effects of GNE-9278 on the conformational and dynamic landscapes of the GluN1/GluN2D receptor to observe how the binding of an allosteric modulator to the pre-transmembrane region alters allosteric communication between the transmembrane and amino-terminal domains. We found that GNE-9278 induces conformational changes far from its site of action by stabilizing the coupled state of the GluN2D amino-terminal domain. This suggests that GNE-9278 demonstrates long-range bottom-up allostery in addition to a localized effect at the transmembrane segments. The conformational landscape of the GluN1/GluN2D receptor in the presence of GNE-9278 resembles that observed for the GluN1/GluN2A receptor, suggesting a similarity in allosteric coupling maintained across different GluN2 subunits and in the presence of small molecule modulators.

## Results

To study the conformational landscape of the extracellular domain of the GluN1/GluN2 NMDA receptor, we utilized smFRET to examine distances throughout the receptor at both the GluN1 and GluN2 subunits. We designed four constructs that allowed us to measure inter-subunit distances of the GluN2 amino-terminal domain and GluN1 transmembrane domain and distances at the GluN1 glycine-binding and GluN2 glutamate-binding domain clefts. We introduced donor-acceptor fluorophores at specific sites within the protein using maleimide chemistry. To ensure specific labeling at our sites of interest, we mutated intrinsic non-disulfide bonded extracellular cysteines (Cys15 and Cys22 in GluN1; Cys231, Cys399, and Cys460 in GluN2A; and Cys24, Cys245, and Cys485 in GluN2D) to serines to form cys-light GluN1 and GluN2 constructs. Additionally, a twin-strep tag was introduced at the C-terminus of GluN1 to facilitate the attachment of the heteromeric receptor to a microscope slide via streptavidin pull-down. For measuring changes near the transmembrane domain (GluN1-TMD/GluN2D), we introduced a cysteine at site Phe554 in the GluN1 pre-M1 segment (Figure 1A). To investigate changes across the GluN2D glutamate-binding domain cleft (GluN1/GluN2D-ABD), we introduced cysteines at sites Gln528 and Met726 (Figure 1B). For measuring changes across the GluN1 glycine-binding domain cleft (GluN1-ABD/GluN2D), we introduced cysteines at sites Ser507 and Thr701 (Figure 1C). For investigating changes across the GluN2 amino-terminal domain (GluN1/GluN2A-ATD and GluN1/GluN2D-ATD), we introduced a cysteine at site Thr174 in GluN2A and Thr188 in GluN2D (Figure 1D). Electrophysiological characterization of the above constructs demonstrated preserved activation, desensitization, and potentiation function, validating their use in smFRET investigations (see Figure 1).

**Figure 1.**
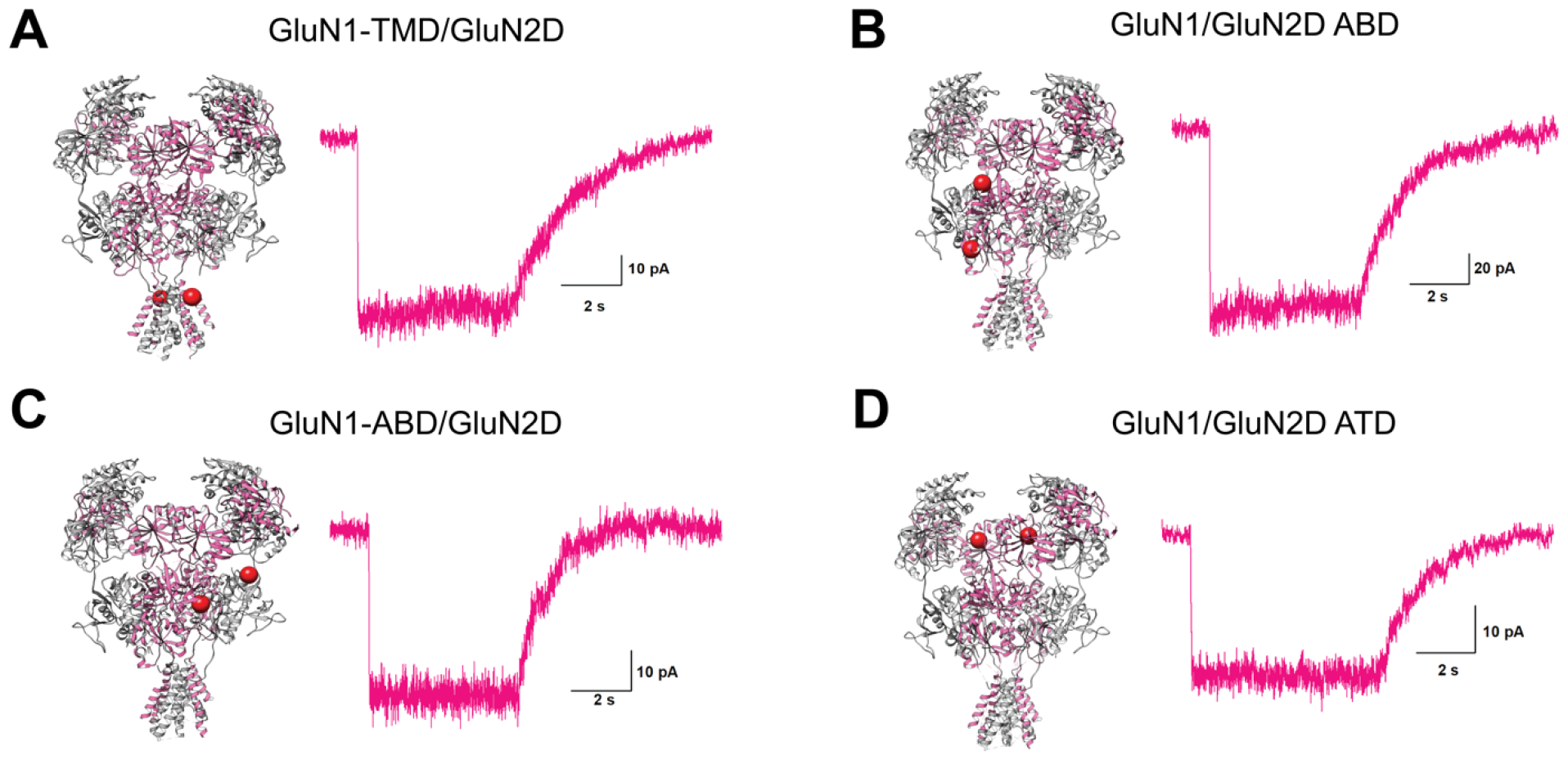
Functional characterization of smFRET constructs. Structure of the GluN1/GluN2D NMDA receptor (PDB ID: 8E96 ^28^) with red spheres representing the location of the fluorophore attachment sites for measuring at **A)** the GluN1 transmembrane domain (F554), **B)** the GluN2 glutamate-binding domain (Q528 and M726), **C)** the GluN1 glycine-binding domain (S507 and T701), and **D)** the GluN2 amino-terminal domain (T188). The traces represent the whole-cell recording for each smFRET construct in the presence of 1 mM glutamate and glycine.

To perform smFRET experiments, HEK293T cells were transfected with GluN1 constructs and either GluN2A or GluN2D constructs (see Methods section for a detailed description). The NMDA receptors were labeled with donor (Alexa 555) and acceptor (Alexa 647) fluorophores. Subsequently, the labeled cells were solubilized and pulled down onto the smFRET slides. A confocal fluorescence microscope was used to visualize fluorophore-labeled receptors and select receptors with one photobleaching FRET for analysis. Donor and acceptor fluorophore emissions were quantified by counting the number of photon detection events in 5 ms time bins, and the FRET efficiency was calculated for each time bin. The resulting FRET traces from individual receptors across multiple independent biological replicates were combined and averaged into cumulative FRET efficiency histograms with standard error bars representing the standard error across histograms from independent samples. Gaussian fitting guided by hidden Markov model-based analysis of individual smFRET traces was used to determine the specific occupied FRET states and quantify the fraction of receptors occupying each FRET state^44, 45^.

### Receptor activation reflected in conformations near pre-M1

Functional studies have determined that GluN1/GluN2A and GluN1/GluN2D receptors vary in open probability, activation time course, and desensitization profiles^17, 18, 19, 20, 21^. Thus, we began by examining conformational rearrangements across the transmembrane segments. We measured distances across the channel pore at the GluN1 subunit pre-M1 segment in the unbound (apo) and glutamate and glycine bound states. While structural studies have not been able to capture the apo conformation of NMDA receptors thus far, we can assess the resting conformations of NMDA receptors with smFRET by applying glycine oxidase to remove contaminating glycine for the unliganded conditions, as previously performed in Durham et al.^46^.

Under apo conditions, we primarily observe a high FRET state of 0.97 which correlates to a FRET distance of approximately 29 Å. This distance agrees with the published cryo-EM structure of the GluN1/GluN2D receptor bound by glycine and competitive antagonist CPP, where the distance between Phe554 on both GluN1 subunits is approximately 31 Å^29^. This finding is consistent with the GluN1 transmembrane segments in the GluN1/GluN2D receptor being tightly packed and likely representing the closed, inactive channel state of the receptor. The spread of states measured at the GluN1 pre-M1 segment demonstrates that the GluN2D-containing receptor is shifted to higher FRET states compared to the GluN1/GluN2A receptor, suggesting an inherent difference at this site under resting conditions, with the GluN1/GluN2D receptor being more tightly packed (Figure 2A)^46, 47^. When both glutamate and glycine are bound to the receptor (using saturating 1 mM concentrations of both ligands), we observe that both GluN1/GluN2D and GluN1/GluN2A shift to occupy lower FRET efficiencies compared to apo conditions. This likely represents the pull on the transmembrane linker that contributes to opening of the channel pore. However, we continue to observe a similar trend of GluN1/GluN2D receptors occupying a higher FRET distribution compared to GluN1/GluN2A receptors, which is consistent with the lower open probabilities of this receptor subtype (Figure 2C).

**Figure 2.**
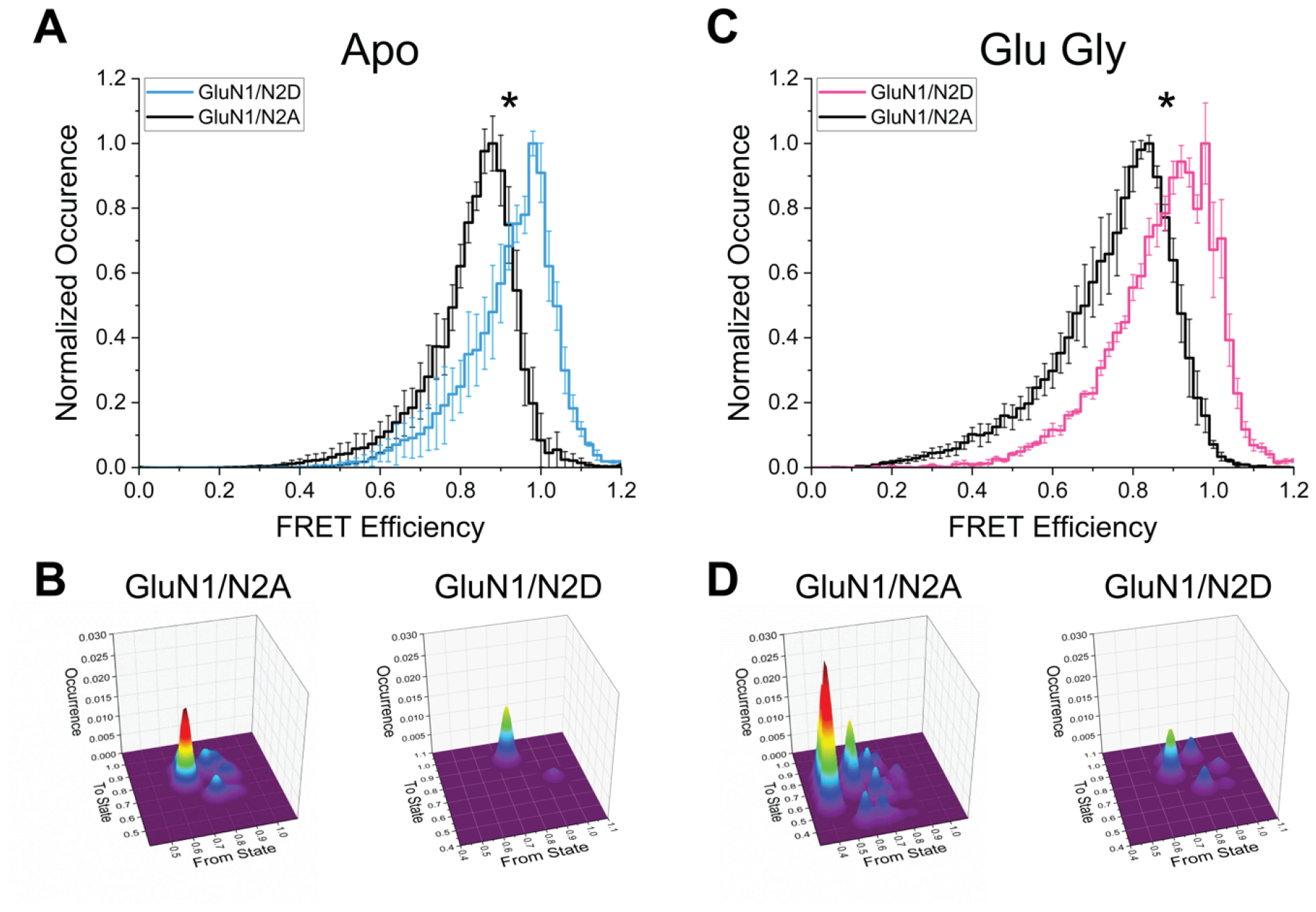
smFRET measurements at the GluN1 transmembrane domain. Cumulative FRET efficiency histograms representing observed FRET efficiencies in the **(A)** apo and **(C)** glutamate and glycine bound conditions obtained from GluN1-TMD/GluN2A (black) and GluN1-TMD/GluN2D (blue for apo; pink for glu/gly) receptors. Additionally, the state transition probability (per 5 ms) maps for the transmembrane domain are shown for the **(B)** apo and **(D)** ligand-bound conditions for both receptor subtypes (GluN1/GluN2A on the left; GluN1/GluN2D on the right). Representative traces are shown in Figure S1. Cumulative histogram values are mean ± SEM of data from at least three independent samples (GluN1/GluN2D N1 TMD apo = 75 mol, 13,196 data points; glu/gly = 74 mol, 14,242 data points). Data for GluN1-TMD/GluN2A are from Litwin et al.^47^. Statistical analysis by two-sample t-test, * *p < 0*.*05*.

The smFRET histograms were fit into states based on Gaussian fitting guided by hidden Markov model-based analysis of the smFRET traces (example traces shown in Supplementary Figure S1). Based on this analysis, we determined that in the unbound condition, the GluN1/GluN2D receptor occupies primarily high FRET states of 0.78 and 0.97 (Table S1). Upon ligand binding, there is an approximately 6% occurrence of a low FRET efficiency of 0.59, which indicates a proportion of receptors exploring longer distances across the transmembrane segments (Table S1). This 6% fraction is consistent with the P_open_ of less than 0.1 reported for GluN1/GluN2D receptors, suggesting that this lower FRET efficiency state reflects channel opening. Additionally, this fraction is significantly smaller than that observed for GluN1/GluN2A receptors, where low FRET efficiencies compose approximately 35%, consistent with the higher P_open_ of approximately 0.5 for this subtype^47^.

The smFRET traces also allow us to investigate conformational dynamics in terms of transition probabilities between states. Transition probability per 5 ms was calculated based on the number of transitions observed in the raw smFRET data while taking into consideration the relative occurrence of the starting state of each smFRET trace. The transition probability maps reveal that the transmembrane segments in both GluN1/GluN2A and GluN1/GluN2D receptors are more dynamic in the fully liganded condition (Figure 2D) compared to the apo condition (Figure 2B). This suggests that the transmembrane segments have increased fluctuations when the receptor is bound by its ligands, allowing for the receptor to populate the low FRET efficiency activated state. Additionally, we observe that the GluN1/GluN2A transmembrane segments are significantly more dynamic than the GluN1/GluN2D transmembrane domain, suggesting a higher free energy barrier for the GluN1/GluN2D transmembrane segments to transition into low FRET states^46, 47^.

This is consistent with the lower open probability of the GluN1/GluN2D receptor and further confirms that conformations arising from low FRET states correlate with pore opening.

### Agonist-induced changes at the glutamate- and glycine-binding domains

To study the agonist-binding domain changes, we measured distances across the bilobed cleft in the GluN1 and GluN2 subunits. The smFRET histograms show that the unbound receptor occupies a broad spread of states at GluN1 and GluN2 agonist-binding clefts in the GluN1/GluN2D receptor (Figure 3A, 3C). For both GluN2 and GluN1 agonist-binding clefts, two-sample t-tests comparing the means of the cumulative FRET efficiency histograms for GluN1/GluN2D and GluN1/GluN2A ^46^ did not return significant p-values. This suggests that the conformational landscape of the agonist-binding domain clefts is similar for both the GluN1/GluN2A and GluN1/GluN2D receptors in the apo condition. Upon binding of glutamate and glycine, the distances across the cleft decrease, and the conformational spread is reduced at both GluN1 and GluN2D subunits (Figure 3B, 3D) (Supplementary Figure S2 and Figure S3). This is consistent with agonist binding stabilizing the agonist-binding domains in a closed cleft conformation. Furthermore, the smFRET histograms for GluN1/GluN2D in the agonist-bound condition at both clefts are not significantly different from those previously observed for GluN1/GluN2A receptors^46, 48, 49^. These studies reveal that the overall conformational landscape of the GluN1 and GluN2 agonist-binding domains are similar, both in the resting and agonist-bound forms between GluN1/GluN2D and GluN1/GluN2A receptors.

**Figure 3.**
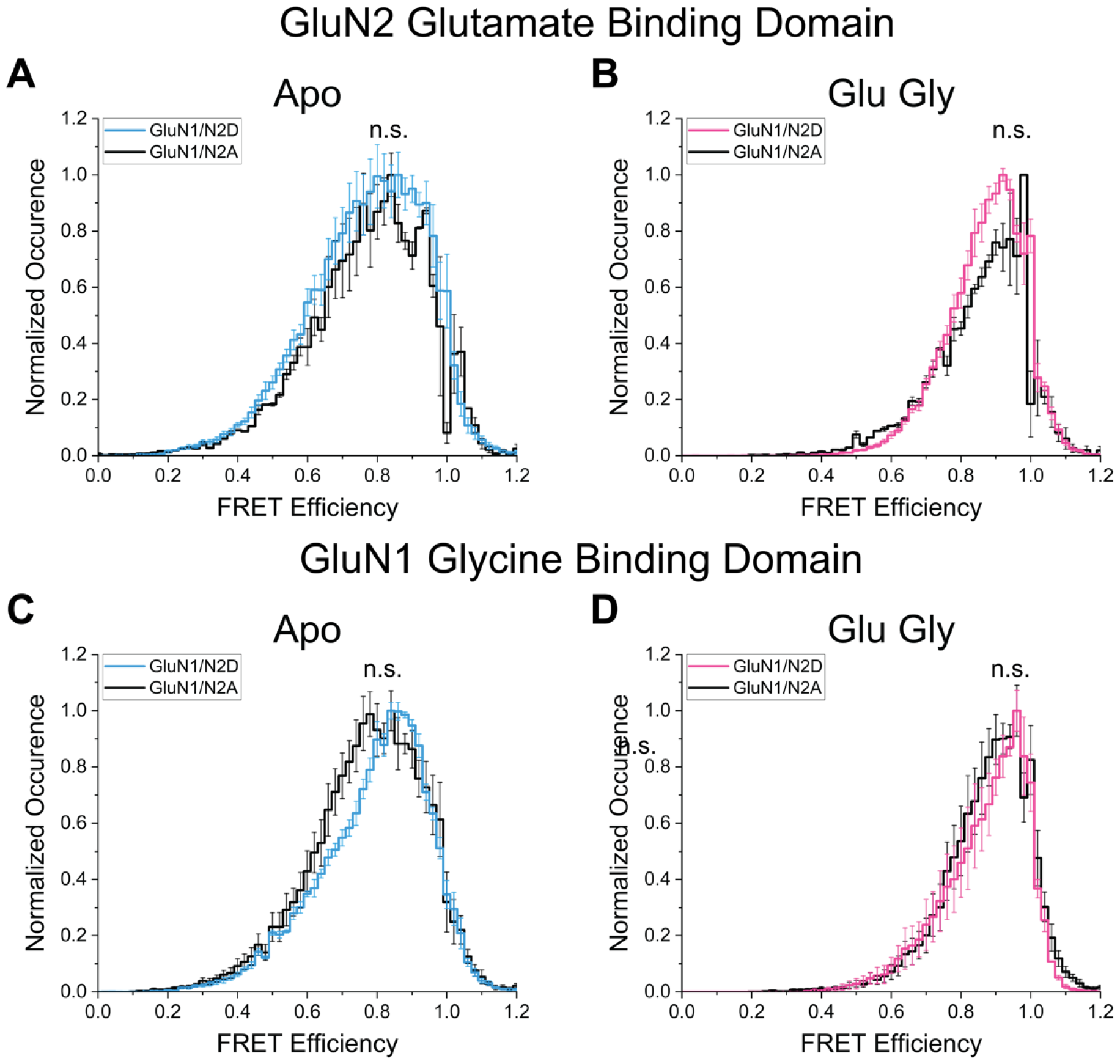
smFRET measurements at the glutamate- and glycine-binding domains. Cumulative FRET efficiency histograms representing observed FRET efficiencies in the **(left, A and C)** apo and **(right, B and D)** glutamate and glycine bound conditions obtained from GluN1/GluN2A (black) and GluN1/GluN2D (blue for apo; pink for glu/gly) receptors at the **(top, A and B)** GluN2 glutamate-binding domain and **(bottom, C and D)** GluN1 glycine-binding domain. Representative traces are shown in Figure S2 and S3. Cumulative histogram values are mean ± SEM of data from at least three independent samples (GluN1/GluN2D N2 ABD apo = 105 mol, 45,097 data points; glu/gly = 110 mol, 48,928 data points; GluN1/GluN2D N1 ABD apo = 85 mol, 42,763 data points; glu/gly = 64 mol, 25,366 data points). Data for GluN1/GluN2A-ABD and GluN1-ABD/GluN2A are from Durham et al.^46^. Statistical analysis by two-sample t-test, * *p < 0*.*05*.

### GluN2-dependent conformations of the amino-terminal domain

Amino-terminal domain conformations dictate differences in receptor activation^22, 50^. Decoupling of the amino-terminal domain upon antagonist or inhibitor binding correlates with lower activity of GluN1/GluN2A and GluN1/GluN2B receptors^29, 51^. To test if a similar shift in conformational landscape underlies lower activity of the GluN1/GluN2D receptor, we studied distances across the GluN2 subunit amino-terminal domain in GluN2D- and GluN2A-containing receptors. The smFRET data reveal a significant trend of lower FRET efficiencies in the apo and agonist-bound conditions for GluN1/GluN2D receptors relative to GluN1/GluN2A receptors (Figure 4A, 4C). This suggests that the GluN2D-containing receptors exhibit more significant decoupling of the amino-terminal domain than GluN2A-containing receptors.

**Figure 4.**
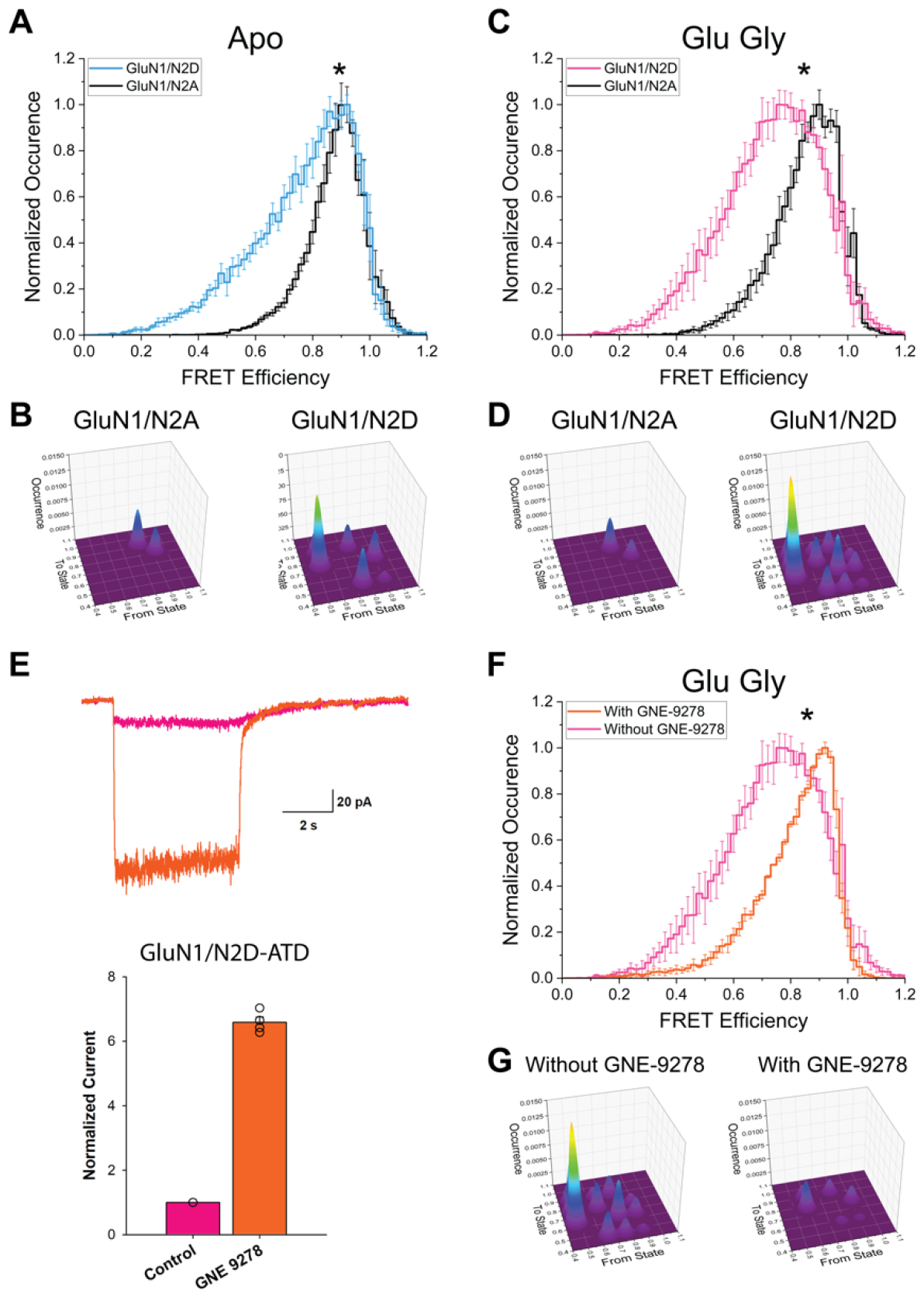
smFRET measurements at the GluN2 amino-terminal domain. Cumulative FRET efficiency histograms representing observed FRET efficiencies in the **(A)** apo, **(C)** glutamate and glycine bound conditions obtained from GluN1/GluN2A-ATD (black) and GluN1/GluN2D-ATD (blue for apo; pink for glu/gly). **(F)** Cumulative FRET efficiency histogram from the glutamate, glycine, and GNE-9278 bound GluN1/GluN2D-ATD (orange) compared to glutamate and glycine bound only form **(C, pink)**. Additionally, the state transition probability (per 5 ms) maps for the amino-terminal domain are shown for the **(B)** apo, **(D)** ligand-bound, and **(G)** ligand and modulator-bound conditions. **(E)** Trace obtained from whole-cell recordings of the GluN1/GluN2D-ATD smFRET construct in the presence of glutamate and glycine (pink) and glutamate, glycine, and GNE-9278 (orange). The bar graph represents the peak amplitude from at least 3 different measurements normalized against the control (glu/gly only, no modulator). Representative traces are shown in Figure S3. Cumulative histogram values are mean ± SEM of data from at least three independent samples (GluN1/GluN2D N2 ATD apo = 83 mol, 34,694 data points; glu/gly = 97 mol, 40,095 data points; glu/gly/GNE-9278 = 80 mol, 31,783 data points; GluN1/GluN2A N2 ATD apo = 91 mol, 40,309 data points; glu/gly = 81 mol, 44,300 data points). Statistical analysis by two-sample t-test, * *p < 0*.*05*.

The smFRET histograms were fit into states based on Gaussian fitting guided by hidden Markov model-based analysis of the smFRET traces (example traces shown in Supplementary Figures S4 and S5). While both GluN2D- and GluN2A-containing receptors occupy similar high FRET efficiency states, GluN2D-containing receptors also populate low FRET efficiency states. This shift towards lower FRET states results in a lower occurrence of the coupled high FRET states. In the glutamate-glycine bound state, the fraction of the high FRET conformation is approximately 10% for the GluN1/GluN2D receptor and 65% for the GluN1/GluN2A receptor (Table S1). These fractions correlate to the P_open_ of less than 0.1 for GluN1/GluN2D receptors and approximately 0.5 for GluN1/GluN2A receptors. Additionally, the 10% occurrence of the super-compact conformation at the GluN2D amino-terminal domain is similar to the 6% occurrence of the low FRET state at the pre-M1 site of GluN1/GluN2D receptors (Table S1)^47^. Given that decoupling at the GluN2 amino-terminal domain has been linked to channel inhibition of GluN1/GluN2A and GluN1/GluN2B receptors and that the fraction of GluN1/GluN2D and GluN1/GluN2A receptors occupying the super-compact state correlates with their respective P_open_, we can conclude that the super-compact conformation may represent the activated state of these receptor.

In addition to the conformational spread, we also investigated the conformational dynamics of the GluN2 amino-terminal domain in terms of transition probabilities between states. Transition probabilities were calculated per 5 ms, as the raw smFRET data was binned to 5 ms. These data show that the GluN2D amino-terminal domain is more dynamic relative to GluN2A in both apo and ligand-bound conditions. In particular, the low FRET states (0.46 and 0.65; Table S1), which are populated by the GluN2D amino-terminal domain upon agonist binding, show many transitions, indicating that these conformations are highly dynamic (Figure 4B and 4D).

### A bottom-up allosteric pathway is revealed through the mechanism of the allosteric modulator GNE-9278

While these studies illustrate the top-down allosteric pathway from the amino-terminal domain to the transmembrane segments, relatively few studies demonstrate a bottom-up allosteric pathway^52^. Several positive allosteric modulators, such as GNE-9278, have been shown to bind to the GluN1 subunit pre-M1 segment. The assumption is that the mechanism of potentiation is primarily through localized changes at the transmembrane segments^39^. It remains unknown if there are long-range conformational alterations. smFRET data at the GluN2 amino-terminal domain in GluN1/GluN2D in the presence of saturating 1 mM glutamate and glycine and 30 μM GNE-9278 show a shift towards higher FRET efficiencies, hence favoring a more coupled state of this domain (Figure 4F). The occurrence of this coupled state is approximately 53%, which correlates to the 6-fold increase in activation observed for the GluN1/GluN2D receptors in the presence of GNE-9278 (Figure 4E; Table S1). Additionally, we observe a decrease in conformational dynamics, suggesting stabilization of the coupled amino-terminal domain conformation upon GNE-9278 binding (Figure 4G). These data show that allosteric communication between the amino-terminal and transmembrane domains is conserved even in the modulatory mechanism of positive allosteric modulators and, more importantly, that the pathway is bidirectional.

## Discussion

The smFRET methodology allows us to explore conformational landscapes in conjunction with protein dynamics at specific sites of interest within a protein guided by existing cryo-EM structures. In turn, this technique allows us to correlate conformations to functional states^53, 54, 55^. Here, we utilize this methodology to study the GluN2 subunit-mediated conformational control of NMDA receptors that correlates with functional differences between receptor subtypes. Our results reveal the occupancy of several common conformational classes between the GluN1/GluN2A and GluN1/GluN2D receptors. However, the population shift in conformation occurrences correlates with the GluN2 subtype-specific functional properties.

The similarity in the conformations occupied at the agonist-binding domains of the GluN1/GluN2A and GluN1/GluN2D receptors suggests that the agonist-binding domains primarily act as a lever switching the receptor from resting to active and desensitized states. The agonist-binding domains do not play a direct role in differences in receptor gating observed between the two subtypes. Furthermore, the lack of differences between GluN1/GluN2A and GluN1/GluN2D receptors at this domain suggests that variations in glutamate and glycine affinity arise from local differences at the agonist-binding site and not due to global cleft closure conformational changes. Such local differences in the side chains and binding site backbone were previously seen in the x-ray structures of the isolated agonist-binding domains of the GluN2A and GluN2D subunits^25, 43, 56^. These structures did not show significant conformational differences, which is consistent with our data on the full-length receptor.

Conformational differences between GluN1/GluN2A and GluN1/GluN2D are primarily seen across the pre-M1 transmembrane segments and at the GluN2 amino-terminal domain. The GluN2 amino-terminal domain of GluN1/GluN2A can be classified into compact and super-compact conformations under agonist-bound conditions. Our data show that the GluN2 amino-terminal domain in GluN1/GluN2D receptors also populates these states, in addition to splayed and super-splayed conformations. The GluN1/GluN2D receptors, which have a low P_open_, demonstrate a shift towards the decoupled splayed and super-splayed conformations relative to the GluN1/GluN2A receptors, which have a high P_open_ and populate the compact and super-compact states to a higher degree (Figure 5). Additionally, in the presence of positive allosteric modulator GNE-9278, the GluN2 amino-terminal domain in GluN1/GluN2D receptors no longer populates the super-splayed state, resulting in a higher occurrence of the super-compact state (Figure 5). This suggests that the compact and super-compact conformations are pre-active and active states, while the splayed and super-splayed conformations represent inactive states.

**Figure 5.**
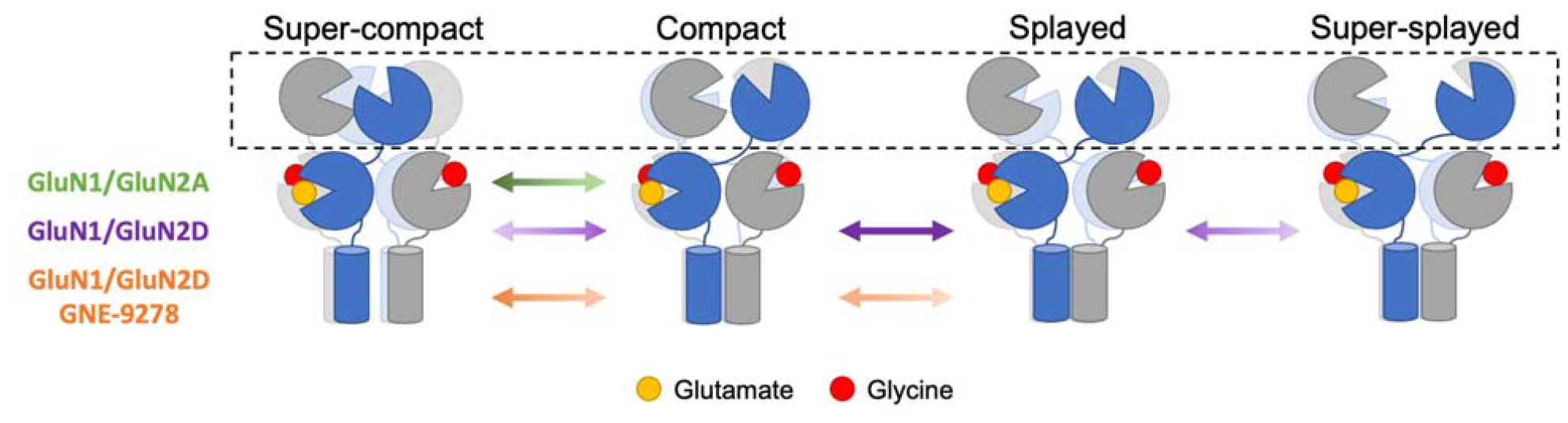
Schematic illustration of the identified conformational states at the GluN2 amino-terminal domain. Cartoon representations of the super-compact, compact, splayed, and super-splayed states that were observed at the GluN2 amino-terminal domain (contained within the dotted box) in agonist-bound GluN1/GluN2A (green), agonist-bound GluN1/GluN2D (purple), and agonist- and GNE-9278-bound GluN1/GluN2D (orange) receptors. GluN1 subunits are shown in gray, and GluN2 subunits are shown in blue. Agonists glutamate and glycine are shown as yellow and red circles, respectively. Arrows represent the relative occurrence of each conformational state, with darker intensity of color indicating higher occurrence while lighter intensity of color indicates lower occurrence. The relative occurrence of each state is listed in Table S1.

While recent cryo-EM structures of GluN1/GluN2D show the coupled conformations (compact and super-compact), the decoupled conformations (splayed and super-splayed) have not been observed^28, 29^. This is most likely due to the highly dynamic nature of these decoupled states, as observed in our smFRET data, which makes them difficult to classify. It is also interesting to note that our smFRET data measuring distances across the GluN2 amino-terminal domain differ from those recently reported for the GluN1 amino-terminal domain^33^. The smFRET investigations measuring distances across the GluN1 amino-terminal domain indicate that the super-compact conformation is inactive, while the compact conformation at this domain is active. These differences indicate that the GluN1 and GluN2 subunits occupy distinct conformations. Such differences were previously observed in cryo-EM structures of the GluN1/GluN2C receptor, which display large splaying at the GluN2 subunits and more minor conformational changes at the GluN1 subunits^28, 29^. It is also important to note that the smFRET measurements performed on the GluN1 subunits were acquired at 100 ms bins. A number of the transition dynamics that we observe across the GluN2D subunits are significantly faster than this binning timescale (Supplementary Figures S5), which would result in the averaging out of the states that we observed in our smFRET data that is binned at 5 ms.

Control of receptor activation and desensitization through inter-subunit coupling and decoupling is seen across different subtypes of the ionotropic glutamate receptor family. Our data reveal that coupling of the GluN2 amino-terminal domain is correlated with receptor activation. Similarly, coupling of the amino-terminal domains in delta receptors through stabilization by cerebellin-1 and neurexin-1β allows these receptors to function as ligand-gated ion channels ^6^. While coupling and decoupling occur at the amino-terminal domain in NMDA and delta receptors, these conformational changes occur at the agonist-binding domain dimer interface in AMPA and kainate receptors. In AMPA and kainate receptors, receptor activation is reflected in the coupling of the agonist-binding domain dimer interface, while decoupling of this interface correlates with desensitization^57, 58, 59, 60, 61, 62^. Additionally, the mechanism of extracellular domain-mediated inter-subunit interaction controlling receptor gating is also observed in the modulation of activation by auxiliary subunits and small molecule modulators^63, 64, 65^. However, it is less understood if these inter-subunit interactions are altered by modulators that bind near the transmembrane segments, away from the extracellular domain. Our smFRET data of GNE-9278 bound GluN1/GluN2D show that binding of the modulator at the transmembrane segments alters the conformations and dynamics of the extracellular amino-terminal domain. Interestingly, recent smFRET and cryo-EM investigations reveal that the allosteric inhibitor GYKI-52466, which binds in the transmembrane collar region of AMPA receptors, also produces a conformational shift in the extracellular domain. Specifically, the binding of GYKI-52466 pushes the AMPA receptor toward occupying a decoupled inactive agonist-binding domain structure^66^. These findings suggest that the allosteric communication from the pre-transmembrane region up toward the extracellular domains is conserved across different subtypes of ionotropic glutamate receptors and establishes the bi-directionality of this allosteric pathway.

## Materials and Methods

### DNA Construct Design

All extracellular nondisulfide-bonded cysteines in wild-type *Rattus norvegicus* GluN1 and GluN2D (NCBI accession codes: NM017010 (GluN1), L31611 (GluN2D), D13211 (GluN2A)) were mutated to serines (C459 in N1; C231, C399, and C460 in GluN2A; C24, C245, and C485 in GluN2D). Site-directed mutagenesis was used to further modify these cys-light constructs to create the constructs used for smFRET measurements. For the GluN1 constructs, a C-terminal Twin-Strep tag was added to GluN1 (GluN1-strep tag), T701 and S507 were changed to cysteines (GluN1-ABD), and, lastly, F554 of GluN1 was changed to a cysteine (GluN1-TMD). The GluN2A-ABD was formed by changing Q503 and M701 to cysteine, and the GluN2A-ATD was created by changing T174 to a cysteine. All GluN1 and the GluN2A-ABD constructs were previously used and shown to be functional by electrophysiology^46, 47, 48, 49^. For the first GluN2D construct, Q528 and M726 of GluN2D were changed to cysteines, and a C-terminal His tag was introduced (GluN2D-ABD). To form the GluN2D-ATD construct, T188 was changed to a cysteine (GluN2D-ATD). All constructs are contained in the pcDNA3.1 vector with a CMV promoter and have been shown to form functional receptors via electrophysiology (see Figure 1). Additionally, all mutations were verified using Sanger sequencing.

### Cell Culture and Transfection

HEK293T cells (ATCC) were cultured at 37°C and 5% CO_2_ in growth media containing Dulbecco’s Modified Eagle’s Medium (DMEM) (GenDepot) supplemented with 10% Fetal Bovine Serum (FBS) (GenDepot), 1 unit/mL Penicillin/ 1 ng/mL Streptomycin (P/S) (Sigma-Aldrich), and 25 mg plasmocin (Invivogen). Before transfection, the cells were cultured for approximately 3-20 passages. At ∼50-70% confluency, the cells were transiently transfected using jetPRIME reagent (Polyplus) with 10 μg of DNA per 10 cm dish in a 1:3 μg ratio of GluN1 to GluN2 DNA. For experiments measuring the GluN2 ATD, cells were transfected with 2.5 μg of GluN1-strep tag and 7.5 μg of GluN2(A or D)-ATD. For experiments measuring the GluN2 ABD, cells were transfected with 2.5 μg of GluN1-strep tag and 7.5 μg of GluN2(A or D)-ABD. For experiments measuring the GluN1 ABD, cells were transfected with 1.25 μg of GluN1-strep tag, 1.25 μg of GluN1-ABD, and 7.5 μg of GluN2(A or D). For experiments measuring the GluN1 TMD, cells were transfected with 2.5 μg GluN1-TMD and 7.5 μg GluN2(A or D). Per smFRET experiment, two 10 cm dishes were transfected. After four hours of transfection, the jetPRIME reagent was replaced with growth media containing 30 μM of GluN1 antagonist DCKA (Abcam) and 300 μM of GluN2 antagonist APV (Abcam) and incubated overnight.

The same procedure was followed for electrophysiology experiments with the following exceptions. At 40-50% confluency, HEK293T cells in 35 mm dishes were transfected using Lipofectamine 2000 (Invitrogen) in optiMEM (Thermo Fisher) and co-transfected with GFP. 30 μM of DCKA and 300 μM of APV were present during and after transfection.

### Electrophysiology

Whole-cell patch-clamp recordings were performed 24-48 hours after transfection using fire-polished borosilicate glass pipettes with 3-5 Ohm resistance, filled with internal solution (135 mM CsF, 33 mM CsOH, 2 mM MgCl2, 1 mM CaCl2, 11 mM EGTA, and 10 mM HEPES, adjusted to pH 7.4 with CsOH). The external solution (140 mM NaCl, 2.9 mM KCl, 1 mM CaCl_2_, and 10 mM HEPES, adjusted to pH 7.4 with NaOH) was locally applied to lifted cells using a stepper motor system (Warner Instruments) in the presence of 1 mM ligands (glutamate and glycine) and in the presence or absence of 30 μM GNE-9278 (Tocris Bioscience).

### smFRET Sample Preparation

Sample preparation was performed 24 hours following transfection. The cells were harvested from the two 10 cm dishes and centrifuged. The cells were then washed in 3 mL of extracellular buffer (ECB) (135 mM NaCl, 3 mM KCl, 2 mM CaCl_2_, 20 mM glucose, and 20 mM HEPES; pH 7.4). The NMDA receptors were labeled using 600 nM Alexa 555-C2-maleimide (Invitrogen) and 2.4 μM Alexa 647-C2-maleimide (Invitrogen) in 3 mL of ECB buffer. To allow the fluorophore dyes to bind to the cysteine residues on the NMDA receptors, the cells were incubated in the dark for 30 minutes at room temperature. Following incubation, the cells were washed with ECB three times and resuspended in 2 mL of Solubilization Buffer (1% Lauryl Maltose Neopentyl Glycol (Anatrace), 2 mM Cholesterol Hydrogen Succinate (MP Biomedicals), and protease inhibitors from Pierce Protease Inhibitor Mini Tablets (ThermoFisher Scientific) in 1X PBS). The sample was nutated at 4°C for 1 hour. Cells were then spun for 1 hour at 4°C in a Beckman Ultracentrifuge using a TLA 100.3 rotor at 44,000 rpm (100,000 rcf). The supernatant was collected and stored on ice before use.

### smFRET Slide Preparation

Bath sonication was used to clean microscope cover glasses in a solution of 5% Liquinox phosphate-free detergent (Alconox Inc.) followed by sonication in acetone. The slides were incubated at 70°C in TI-1 solution (4.3% NH_4_OH and 4.3% H_2_O_2_) for 5 minutes. The slides were washed and dried before being subjected to plasma cleaning using a Harrick Plasma PDC-32G Plasma Cleaner. The slides were aminosilanized using a 1:40 Vectabond (Vector Laboratories) to acetone solution. Cleaned and Vectabond-treated slides were stored under vacuum.

The day before each experiment, an overnight PEG solution (0.25% w/w 5 kDa biotin-terminated PEG (NOF Corp.) and 25% w/w 5 kDa mPEG succinimidyl carbonate (Laysan Bio Inc.) in 10 mM sodium bicarbonate) was applied. On the day of each experiment, the slides were treated with a short-chain PEG solution (25 mM short-chain 333 Da MS(PEG)4 Methyl-PEG-NHS-Ester Reagent (Thermo Scientific) in 0.1 M sodium bicarbonate) and incubated for 2 to 3 hours at room temperature. A physical chamber was constructed on top of the slides using adhesive Hybriwell chambers (Grace Bio-labs) and press-fit tubing connectors (Fisher Scientific). 36 μL of streptavidin solution (0.2 mg/mL Streptavidin (Invitrogen) in 1X Imaging Buffer (1 mM nDodecyl-beta-D-maltoside (Chem-Impex Int’l Inc.) and 0.2 mM Cholesteryl Hydrogen Succinate (MP Biomedicals) in 1X PBS)) was loaded into the slide chamber and allowed to incubate for 10 minutes at room temperature. When the molecule of interest did not contain a Twin-Strep Tag, 10 nM of biotinylated goat Anti-Mouse IgG (H+L) secondary antibody (Jackson ImmunoResearch Laboratories, Inc., cat. #115-065-003) was loaded into the chamber and incubated for 20 minutes. The chamber was flushed using buffer to remove unbound antibodies before applying 10 nM of anti-GluN1 mouse monoclonal primary antibody (Synaptic Systems, cat. #114 011). Antibody treatment was not necessary when the molecule of interest contained a genetically encoded Twin-Strep Tag. The supernatant prepared in the smFRET Sample Preparation section above was either directly applied to the slide or diluted between 1:2 and 1:5 in 1X Imaging Buffer before application. The slides were incubated for 20 minutes at 4°C. The slide chambers were washed with reactive oxygen species (ROXS) scavenging solution (3.3% w/w glucose (Sigma-Aldrich), 3 units/mL pyranose oxidase (Sigma-Aldrich), 0.001% w/w catalase (Sigma-Aldrich), 1 mM ascorbic acid (Sigma-Aldrich), and 1 mM methyl viologen (Sigma-Aldrich) in 1X imaging Buffer, pH 7.5; plus 1 mM glutamate and/or 1 mM glycine and/or 12 ug/mL glycine oxidase (Biovision) and/or 30 μM GNE-9278 as needed to create the appropriate experimental conditions). The slides were then imaged.

### smFRET Data Collection and Analysis

A PicoQuant MicroTime 200 Fluorescence Lifetime Microscope was used to acquire the smFRET data. The fluorophores were excited using 532 nm (LDH-D-TA-530; Picoquant) and 637 nm (LDH-D-C-640; Picoquant) lasers operated in pulsed interleaved excitation (PIE) mode with an 80 MHz pulse rate. The slides were mounted on an x-y-z piezo stage (P-733.2CD; Physik Instrumente), and single molecules were observed through an oil-immersed 100x objective. 550 nm and 650 nm emission filters were used to collect the fluorescence emission.

Donor and acceptor fluorophore intensities for all individual molecules were measured by the photon detection counts per 5 ms time bin, and the corresponding FRET efficiency for each time bin was calculated. To ensure that the final traces were obtained from molecules with a single FRET pair attached, molecules were screened for traces showing anticorrelation between the donor and acceptor intensities as well as a single photobleaching step in the donor and acceptor channels. Photoblinking events were removed from the selected smFRET traces. The total number of molecules and total number of data points (binned to 5 ms) included in the final analysis for each condition was as follows: **GluN2D N2 ATD** apo = 83 molecules, 34,694 data points, glu/gly = 97 molecules, 40,095 data points, glu/gly/GNE = 80 molecules, 31,783 data points; **GluN2D N2 ABD** apo = 105 molecules, 45,097 data points, glu/gly = 110 molecules, 48,928 data points; **GluN2D N1 ABD** apo = 85 molecules, 42,763 data points, glu/gly = 64 molecules, 25,366 data points; **GluN2D N1 TMD** apo = 75 molecules, 13,196 data points, glu/gly = 74 molecules, 14,242 data points; **GluN2A N2 ATD** apo = 91 molecules, 40,309 data points, glu/gly = 81 molecules, 44,300 data points. All smFRET data for the GluN2A N2 ABD, GluN2A N1 ABD, and GluN2A N1 TMD under apo and glu/gly conditions was previously published^46, 47^.

After correcting all individual traces for background signal, cross-talk between donor and acceptor channels, differences in quantum yield, and detector efficiency, FRET efficiency over time for each molecule was calculated using the intensity of the acceptor and donor fluorophores with the Förster equation. The individual molecules were pooled for each measurement site and ligand condition to create cumulative histograms of the FRET efficiency distribution in MATLAB (MathWorks). The histograms provide a visual representation of the fractional occurrence of various smFRET efficiency states. The total number of molecules collected for final analysis was obtained from 2 to 6 days of experiments. The mean and SEM of each measurement site and condition were determined from the data across the different days and significance was determined by comparing the means using a two-sample t-test. Datasets with p-values <0.05 are considered to be significant. Using Origin software (OriginLab), Gaussian fitting guided by Hidden Markov model-based analysis of smFRET traces was used to reveal underlying conformational states that comprise the overall FRET efficiency distribution and transition probabilities^44, 45^. A summary of the preferred FRET efficiency states for all constructs and conditions alongside calculated FRET distances are shown in Table S1. A detailed protocol for smFRET data collection and analysis has been detailed in book chapters by Litwin et al. and Paudyal et al.^53, 55^.

To ensure that our data acquisition and analysis were not biased, one set of data for the amino-terminal domain smFRET was obtained blind for each investigated condition. The samples were prepared and unidentified by one lab member and collected and analyzed by a second without knowledge of the conditions. We observed that this set was statistically similar to the data obtained without blinding.

## Acknowledgments

This project was supported by a training fellowship from the Gulf Coast Consortia on the Houston Area Molecular Biophysics Program (Grant No. T32 GM008280) to Paula Bender and National Institutes of Health Grant (R35GM122528) to Vasanthi Jayaraman. The authors declare no competing financial interests.

## Author Contributions

Paula A. Bender performed the smFRET measurements, analyzed the smFRET data, and wrote and edited the manuscript. Elisa Carrillo performed the electrophysiology measurements and analyzed the electrophysiology data. Subhajit Chakraborty, Ryan J. Durham, and Vladimir Berka performed the smFRET experiments and analyzed the smFRET data. Vasanthi Jayaraman designed the research, analyzed the data, and wrote and edited the manuscript.

