## Supplementary material for "Bi-directional allosteric pathway in NMDA receptor activation and modulation": Supplemetal Figure and Table

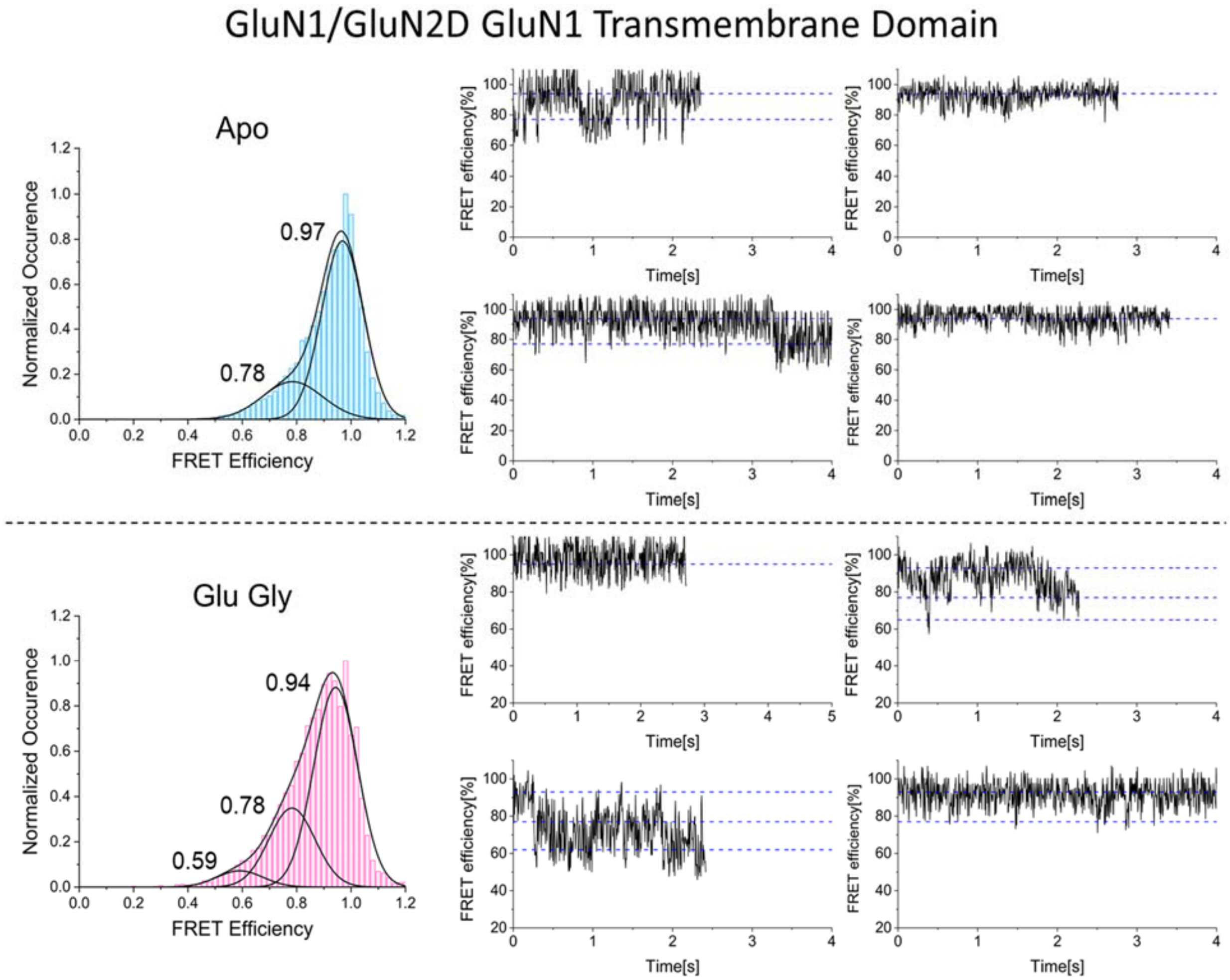


**Figure S1. Gaussian fitting and representative smFRET efficiency traces for the GluN1 transmembrane domain in the GluN1/GluN2D receptor.** Conformational states were predicted using hidden Markov model-based analysis of individual smFRET efficiency traces. The predicted states were used to inform Gaussian fittings (black) of the cumulative FRET efficiency histograms in the apo (blue) and glutamate and glycine bound (pink) conditions. Representative smFRET efficiency traces for the GluN1 transmembrane domain in the GluN1/GluN2D receptor in the presence of no ligand or glutamate and glycine are shown above.

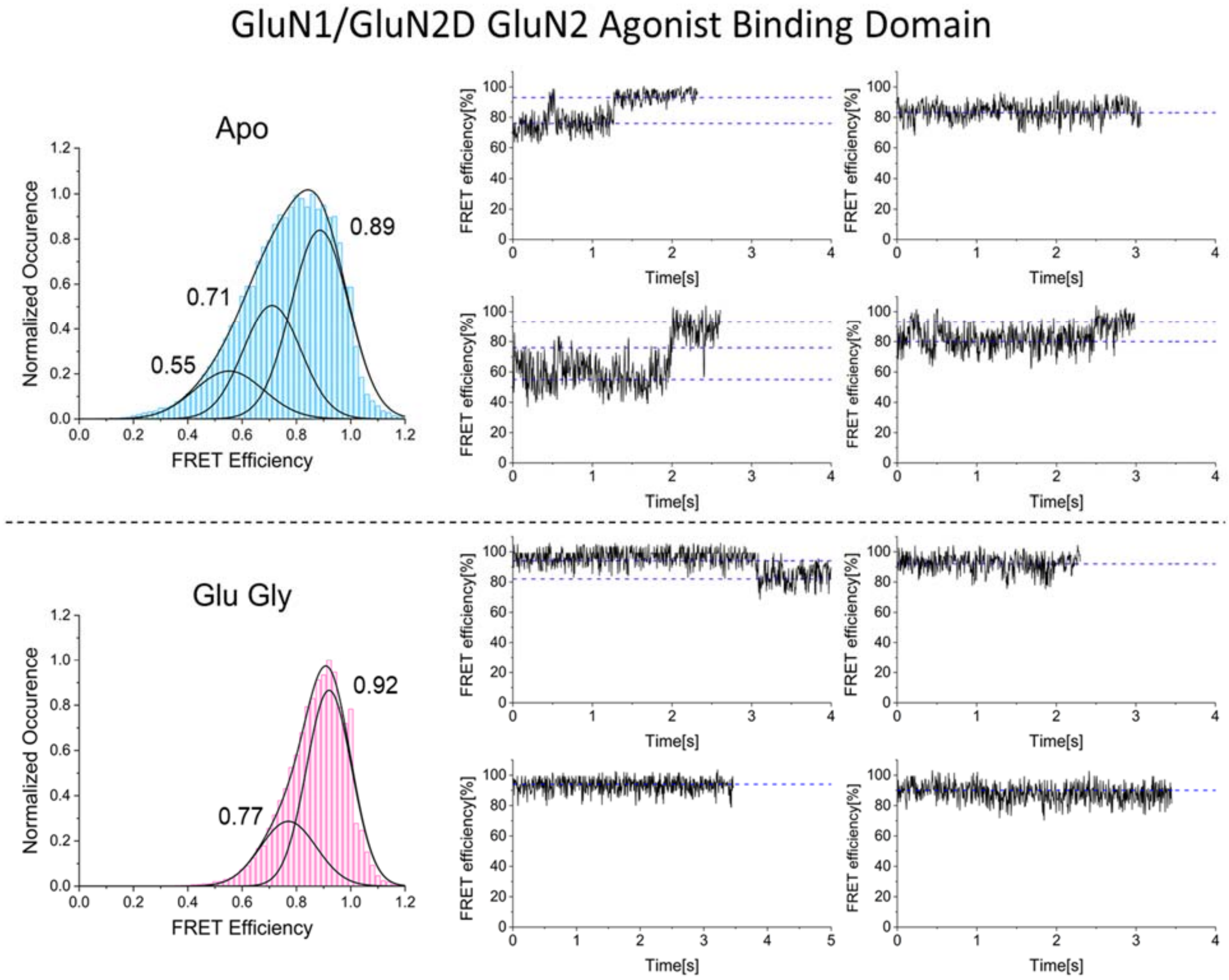


**Figure S2. Gaussian fitting and representative smFRET efficiency traces for the GluN2 agonist binding domain in the GluN1/GluN2D receptor.** Conformational states were predicted using hidden Markov model-based analysis of individual smFRET efficiency traces. The predicted states were used to inform Gaussian fittings (black) of the cumulative FRET efficiency histograms in the apo (blue) and glutamate and glycine bound (pink) conditions. Representative smFRET efficiency traces for the GluN2 agonist binding domain in the GluN1/GluN2D receptor in the presence of no ligand or glutamate and glycine are shown above.

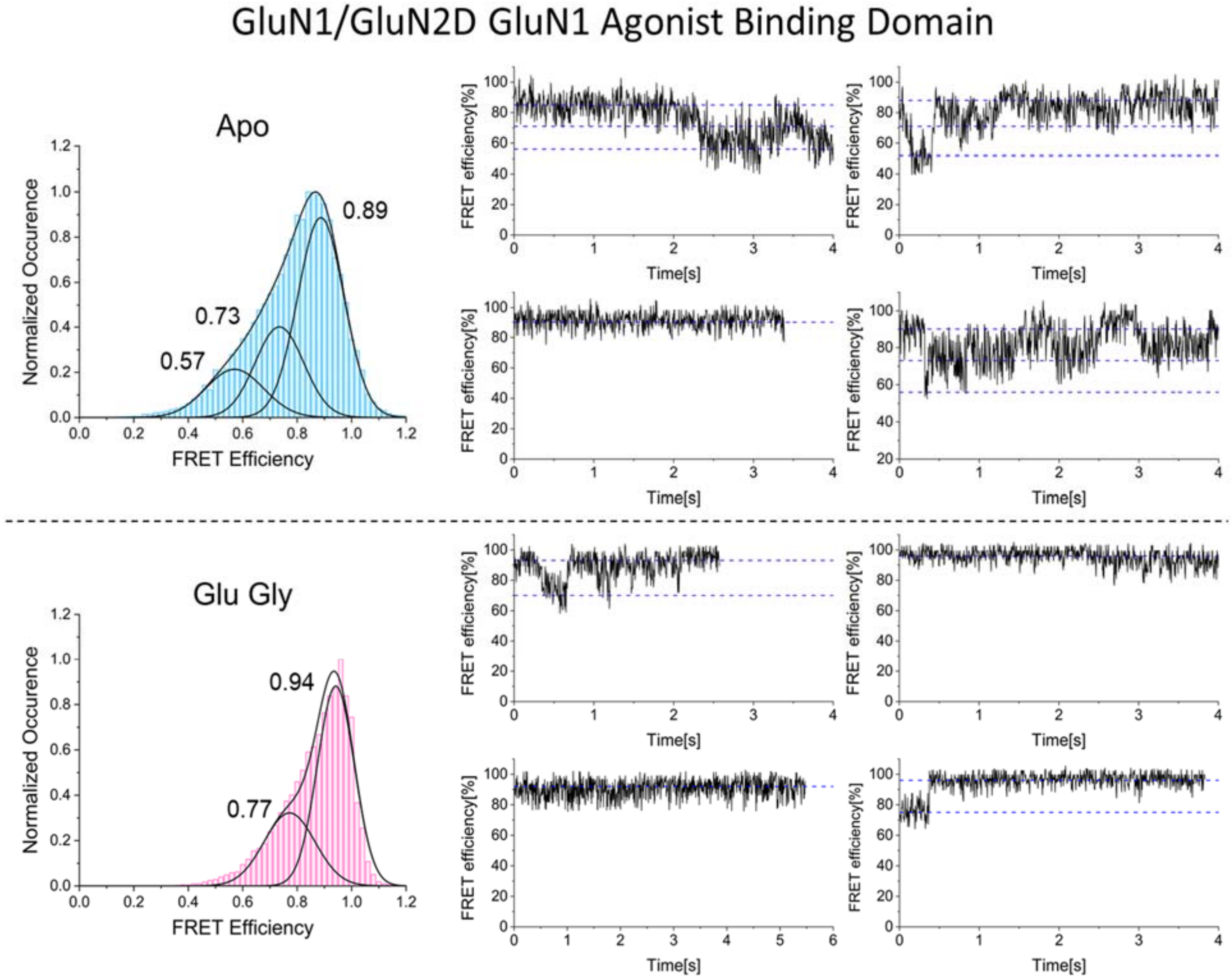


**Figure S3. Gaussian fitting and representative smFRET efficiency traces for the GluN1 agonist binding domain in the GluN1/GluN2D receptor.** Conformational states were predicted using hidden Markov model-based analysis of individual smFRET efficiency traces. The predicted states were used to inform Gaussian fittings (black) of the cumulative FRET efficiency histograms in the apo (blue) and glutamate and glycine bound (pink) conditions. Representative smFRET efficiency traces for the GluN1 agonist binding domain in the GluN1/GluN2D receptor in the presence of no ligand or glutamate and glycine are shown above.

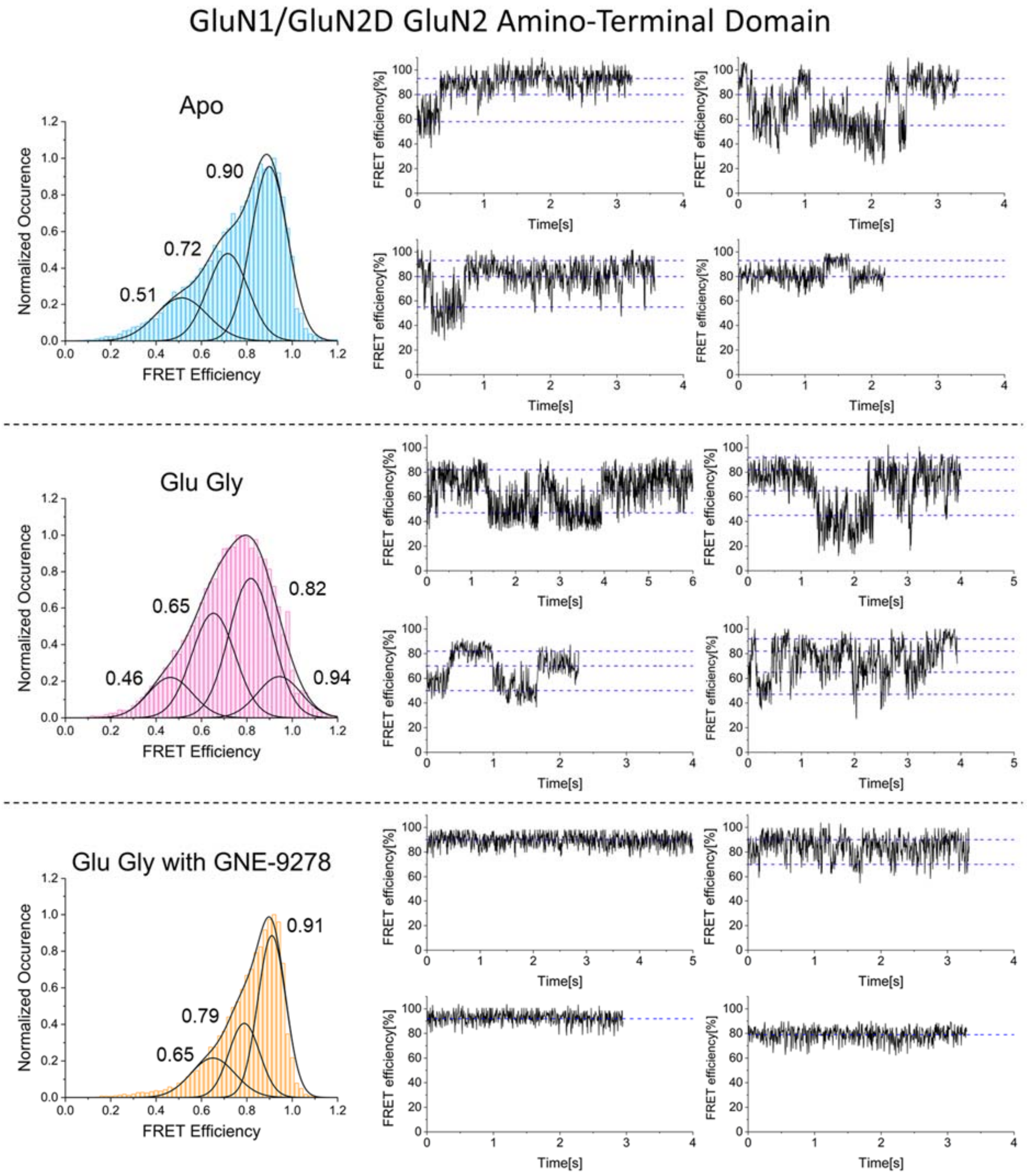


**Figure S4. Gaussian fitting and representative smFRET efficiency traces for the GluN2 amino-terminal domain in the GluN1/GluN2D receptor.** Conformational states were predicted using hidden Markov model-based analysis of individual smFRET efficiency traces. The predicted states were used to inform Gaussian fittings (black) of the cumulative FRET efficiency histograms in the apo (blue), glutamate and glycine bound (pink), and glutamate/glycine/GNE-9278 bound conditions. Representative smFRET efficiency traces for the GluN2 amino-terminal domain in the GluN1/GluN2D receptor in the presence of no ligand, glutamate and glycine, or glutamate and glycine with GNE-9278 are shown above.

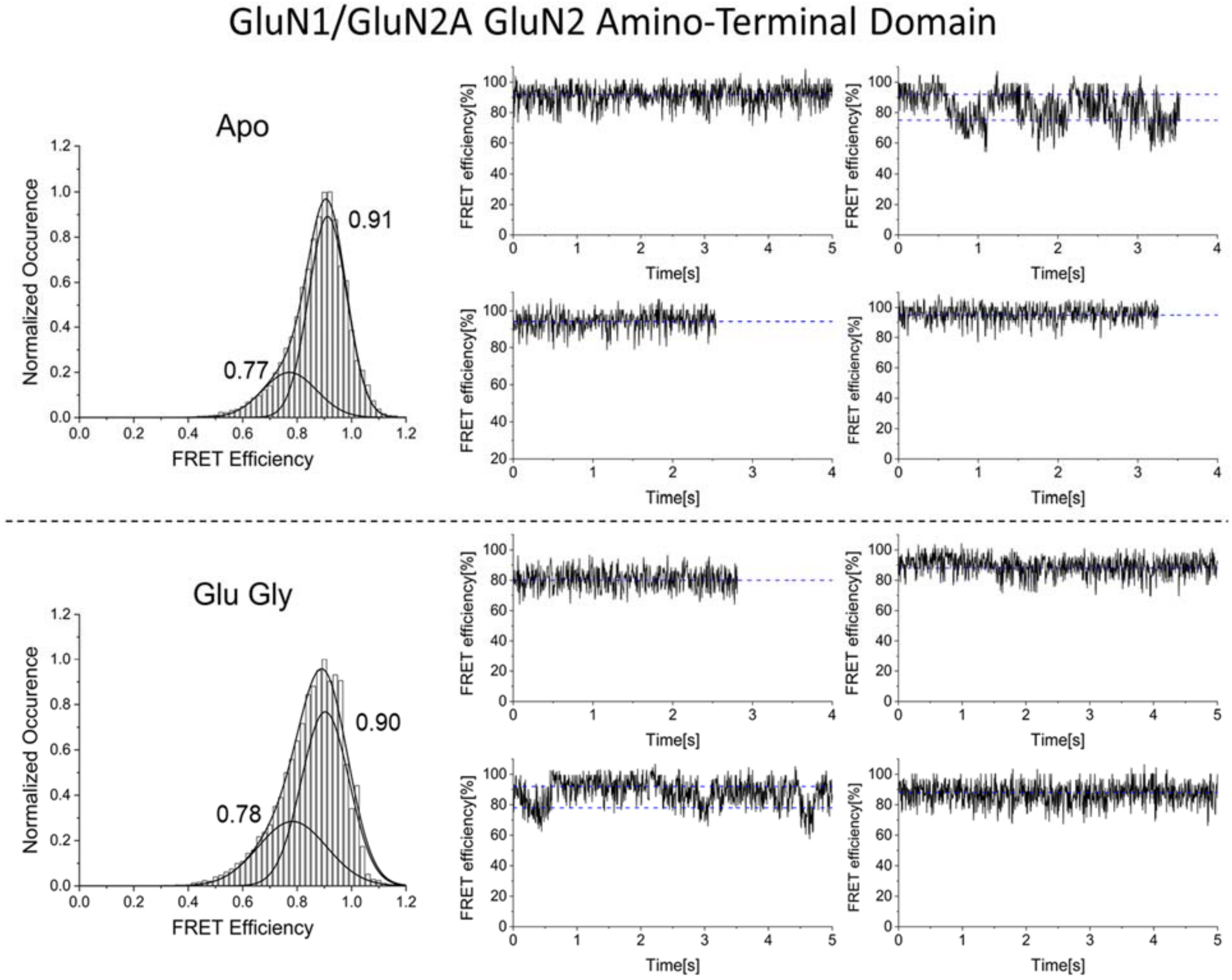


**Figure S5. Gaussian fitting and representative smFRET efficiency traces for the GluN2 amino-terminal domain in the GluN1/GluN2A receptor.** Gaussian fitting and representative smFRET efficiency traces for the GluN2 amino-terminal domain in GluN1/GluN2A. Conformational states were predicted using hidden Markov model-based analysis of individual smFRET efficiency traces. The predicted states were used to inform Gaussian fittings (black) of the cumulative FRET efficiency histograms in the apo and glutamate and glycine bound conditions. Representative smFRET efficiency traces for the GluN2 amino-terminal domain in the GluN1/GluN2A receptor in the presence of no ligand or glutamate and glycine are shown above.

| GluN1 Transmembrane Domain States in GluN1/GluN2D | | | |
|---|---|---|---|
| | FRET Efficiency State | Percent Occurrence | FRET Distance (Å) |
| Apo | 0.78 ± 0.003 | 23.3% | 41.3 ± 0.1 |
| | 0.97 ± 0.02 | 76.7% | 28.6 ± 3.3 |
| Glu Gly | 0.59 ± 0.004 | 5.7% | 48.0 ± 0.1 |
| | 0.78 ± 0.02 | 27.7% | 41.3 ± 0.8 |
| | 0.94 ± 0.005 | 66.6% | 32.2 ± 0.5 |
| **GluN2 Agonist Binding Domain States in GluN1/GluN2D** | | | |
| | FRET Efficiency State | Percent Occurrence | FRET Distance (Å) |
| Apo | 0.55 ± 0.04 | 15.9% | 49.3 ± 1.3 |
| | 0.71 ± 0.03 | 31.6% | 43.9 ± 1.1 |
| | 0.89 ± 0.007 | 52.5% | 36.0 ± 0.4 |
| Glu Gly | 0.77 ± 0.02 | 29.1% | 41.7 ± 0.8 |
| | 0.92 ± 0.004 | 70.9% | 33.9 ± 0.3 |
| **GluN1 Agonist Binding Domain States in GluN1/GluN2D** | | | |
| | FRET Efficiency State | Percent Occurrence | FRET Distance (Å) |
| Apo | 0.57 ± 0.01 | 16.9% | 48.7 ± 0.3 |
| | 0.73 ± 0.01 | 26.8% | 43.2 ± 0.4 |
| | 0.89 ± 0.002 | 56.3% | 36.0 ± 0.1 |
| Glu Gly | 0.77 ± 0.01 | 33.4% | 41.7 ± 0.4 |
| | 0.94 ± 0.003 | 66.6% | 32.2 ± 0.3 |
| **GluN2 Amino-Terminal Domain States in GluN1/N2D** | | | |
| | FRET Efficiency State | Percent Occurrence | FRET Distance (Å) |
| Apo | 0.51 ± 0.01 | 18.4% | 50.7 ± 0.5 |
| | 0.72 ± 0.008 | 29.1% | 43.6 ± 0.3 |
| | 0.90 ± 0.003 | 52.5% | 35.4 ± 0.2 |
| Glu Gly | 0.46 ± 0.02 | 13.0% | 52.4 ± 0.7 |
| | 0.65 ± 0.02 | 33.0% | 46.0 ± 0.7 |
| | 0.82 ± 0.03 | 44.1% | 39.6 ± 1.3 |
| | 0.94 ± 0.05 | 9.9% | 32.2 ± 4.8 |
| Glu Gly GNE-9278 | 0.65 ± 0.02 | 20.5% | 46.0 ± 0.7 |
| | 0.79 ± 0.01 | 26.7% | 40.9 ± 0.4 |
| | 0.91± 0.003 | 52.8% | 34.7 ± 0.2 |
| **GluN2 Amino-Terminal Domain States in GluN1/N2A** | | | |
| | FRET Efficiency State | Percent Occurrence | FRET Distance (Å) |
| Apo | 0.77 ± 0.05 | 24.0% | 41.7 ± 2.0 |
| | 0.91 ± 0.005 | 76.0% | 34.7 ± 0.4 |
| Glu Gly | 0.78 ± 0.03 | 34.7% | 41.3 ± 1.2 |
| | 0.90 ± 0.004 | 65.3% | 35.4 ± 0.3 |

**Table S1. Summary of preferred FRET efficiency states sampled by GluN1/GluN2D and GluN1/GluN2A at various domains under different conditions.** Conformational states were first predicted using hidden Markov-based analysis of individual smFRET traces. Gaussian curves informed by predicted states were then fit to the observed FRET efficiency histograms. The percent occurrence of each state was calculated from the relative area under each Gaussian curve. Distance between the fluorophore labeling pairs for each state was calculated based on the centers of the Gaussian curves using an $R_0$ of 51 Å.
